# Colonization during a key developmental window reveals microbiota-dependent shifts in growth and immunity during undernutrition

**DOI:** 10.1101/2023.07.07.547849

**Authors:** Yadeliz A. Serrano Matos, Jasmine Cano, Hamna Shafiq, Claire Williams, Julee Sunny, Carrie A. Cowardin

## Abstract

Childhood undernutrition is a major global health challenge with devastating lifelong consequences. Linear growth stunting due to undernutrition has been linked to poor outcomes, and mothers who experience stunting are more likely to give birth to stunted children. Murine models that capture the intergenerational and multifactorial nature of undernutrition are critical to understanding the underlying biology of this disorder. Here we report a gnotobiotic mouse model of undernutrition using microbiota from human infants with healthy or stunted growth trajectories. Intergenerational transmission of microbiota from parents to offspring leads to the development of growth and immune features of undernutrition and enteropathy, including reduced linear growth, intestinal villus blunting and accumulation of intraepithelial lymphocytes. In contrast, colonization after weaning reduces sensitivity to detect changes driven by distinct microbial communities. Overall, these results suggest intergenerational colonization is a useful approach with which to investigate microbiota-dependent growth and immunity in early life.

## Introduction

Childhood undernutrition is a formidable global health challenge, contributing to nearly half of all deaths in children under the age of five ^1,2^. The first 1,000 days of a child’s life are particularly critical in determining developmental outcomes, and undernutrition during this time can have devastating consequences ^3,4^. Linear growth stunting (length-for-age Z score ≤ 2 standard deviations below the WHO median) is a major complication of undernutrition impacting 149.2 million children globally in 2020 ^1,5^. Stunted mothers are more likely to give birth to stunted children, leading to a cycle of intergenerational transmission that has proven difficult to disrupt ^6,7^. Indeed, some of the best predictors of attained height in children are the child’s weight and length at birth and mother’s attained height, emphasizing the importance of development *in utero* and in early life ^8–11^.The negative consequences of growth stunting persist into adulthood and include poor cognitive development, reduced educational attainment, and increased risk of metabolic and infectious disease ^4,12^. This syndrome is multifactorial and driven by inadequate nutrition, altered gut microbial communities and intestinal inflammation caused by pervasive pathogen colonization ^13,14^. These combined insults can drive a subclinical syndrome of intestinal epithelial derangement and poor absorptive capacity known as Environmental Enteric Dysfunction (EED), which is prevalent in areas with high rates of undernutrition. EED is thought to limit the efficacy of therapeutic foods by decreasing the absorptive capacity of the small intestine ^15–17^. Hallmark features of EED include epithelial remodeling and immune activation, with increased numbers of small intestinal antibody-producing plasma cells, regulatory and cytotoxic T cells, and reductions in macrophages ^18–20^. Despite measurable progress in reducing stunting due to undernutrition, many of its long term consequences have proven resistant to pharmaceutical and nutritional therapies, highlighting the need to further understand the underlying etiology and mechanisms driving pathology ^21,22^.

The gut microbiome plays a critical role in shaping both local and systemic immunity, and children with undernutrition and EED are known to have altered gut microbial communities ^23–25^. Transplantation of microbes from undernourished human donors to recipient germ-free animals (GF) suggests these community alterations are causally linked to deficits in growth and alterations in metabolism ^25,26^. Murine models have likewise highlighted early life as a critical window in which the immune system develops in reaction to the gastrointestinal microbiome ^27,28^. Indeed, GF mice that are colonized after the weaning period display heightened susceptibility to inflammatory pathologies ^28^. Thus, the timing of colonization can dictate later immune outcomes, with significant implications for the long-term sequelae of undernutrition, many of which can be linked to immune dysfunction. These findings also raise the question of whether immunity differs depending on the composition of the microbiota that is present during critical periods of development.

To address these questions, we designed a murine model of early life undernutrition using human gut microbiota transmitted vertically from dams to offspring. In this “intergenerational” model, GF breeding mice were colonized with microbiota obtained from human infants with healthy or stunted growth trajectories. Offspring born to these mice inherited distinct microbial communities and were weaned onto a nutrient-deficient diet, capturing a critical window of early life development. We compared these mice to animals born GF but colonized directly after the weaning period. We demonstrate that intergenerational colonization with gut microbiota from human infant donors with linear growth stunting leads to significant intestinal immune alterations reminiscent of those seen in human cohorts with EED. These changes arise when animals are born to colonized dams, but not when colonized after weaning. We suggest that this model may serve as a useful tool to delineate the role of specific microbiota-dependent immune changes and their functional consequences during early life undernutrition.

## Results

### Establishing an intergenerational model of undernutrition

In order to determine whether immune developmental outcomes would differ depending on the composition of the gut microbiota during early life, we colonized young (4 week-old) GF C57Bl/6 mice with fecal microbiota sampled from one of four donors: two healthy (height-for-age Z score (HAZ) = 1.12 and 1.74) or two severely stunted (HAZ = −3.35 and −2.33) six month old Malawian infants (**Fig. S1A**) ^23,29^. Mice were fed a micro and macro-nutrient deficient diet composed of foods commonly consumed by the donor population (Malawi-8 or M8 ^23^). In one arm of the experiment, male and female mice were colonized at 4 weeks of age and maintained on the M8 diet until 8 weeks, when microbiota composition and immune phenotypes were assessed. These animals represent the “Post-Weaning” (PW) model of colonization (**Fig. 1A**). In a second arm of the experiment, male mice were similarly colonized and maintained on M8 from 4 to 8 weeks of age. At 8 weeks, age-matched GF females were introduced, breeding pairs were cohoused and maintained thereafter on a nutrient-sufficient diet. Male and female offspring from these breeding pairs were weaned onto the M8 diet from 3 to 8 weeks of age. These animals represent the “Intergenerational” (IG) model of colonization (**Fig. 1A**). To identify phenotypes that differed based on donor growth status, we assessed results by combining data from both stunted donors (SD) and comparing against combined data from both healthy donors (HD).

**Figure 1.**
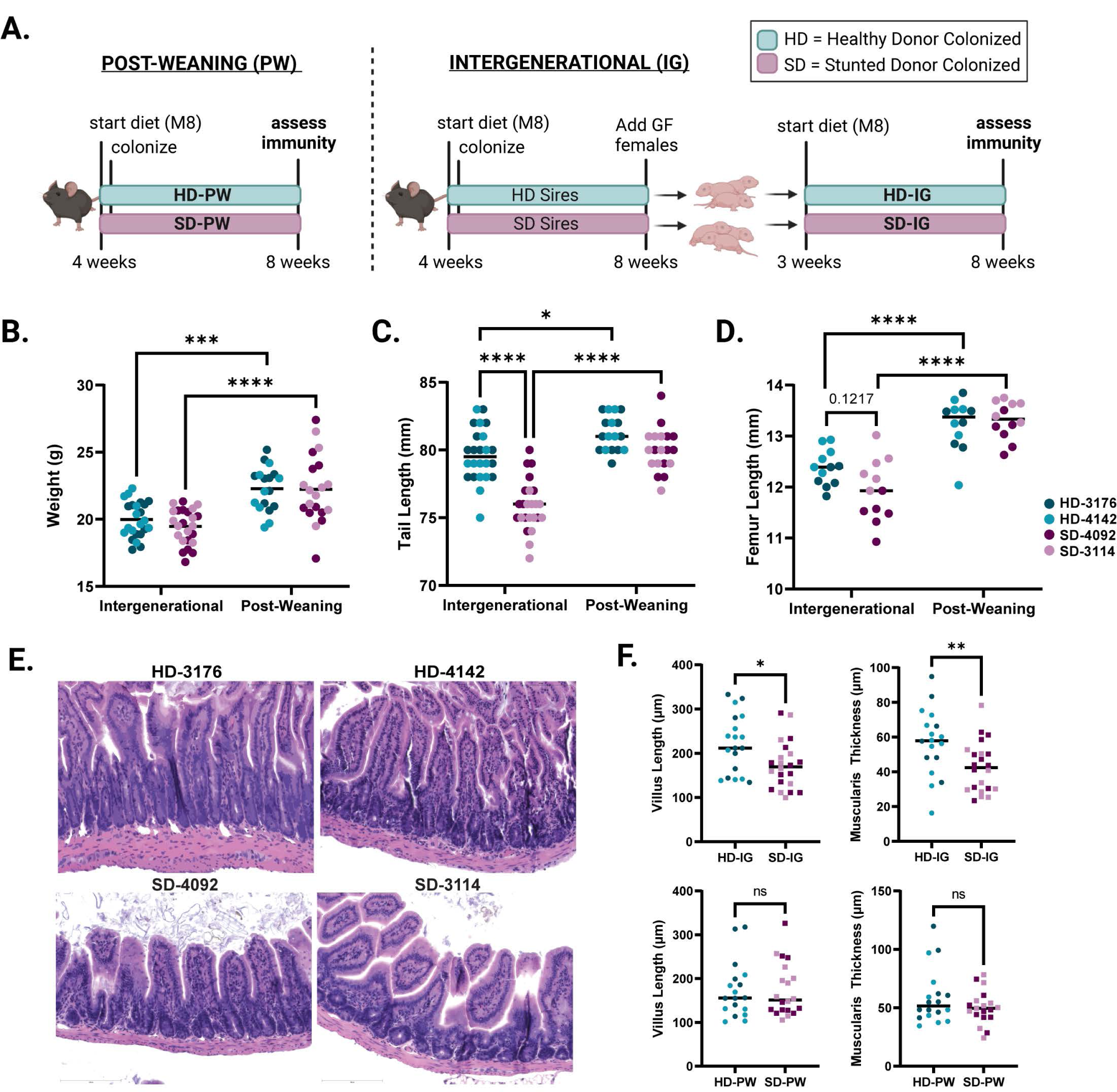
Development of a murine model of intergenerational undernutrition. **(A)** Schematic of experimental design (created with Biorender.com) **(B)** Absolute weight of animals at the time of euthanasia at 8 weeks of age. **(C)** Tail length at 8 weeks of age. **(D)** Femur length at 8 weeks of age. **(E)** Representative histological images of H&E stained ileal tissue from IG mice. **(F)** Quantification of villus length and muscularis thickness in IG and PW mice. Data shown pooled between healthy donor colonized mice (3176 and 4142) and stunted donor colonized mice (4092 and 3114). Each point represents an individual animal. (**B-D**) * p ≤ 0.05, **** p ≤ 0.0001 by Two-Way ANOVA with Šídák’s multiple comparisons test. (**B-C**) n = 24/group [12 per donor] for IG groups, n=18-20/group [8-10 per donor] for PW groups. (**D**) n = 12/group [6 per donor] for all groups in all conditions (**F**) n =18-22/group [7-12 per donor] for IG groups, n = 18-20/group [8-10 per donor] for PW groups. * p ≤ 0.05, ** p ≤ 0.01 by Mann-Whitney U test.

### Intergenerational colonization with SD microbiota reduces linear growth

We next assessed growth phenotypes in both IG and PW animals at 8 weeks of age. We found no difference in overall weight between HD and SD animals colonized either intergenerationally or post-weaning (**Fig. 1B**); however, both groups weighed significantly more when colonized after weaning, possibly due to an additional week of undernutrition in the IG groups. Consistent with these results, we found no significant differences in weight between groups of IG dams and sires after colonization, or between SD-PW and HD-PW mice in the first two weeks after colonization **(Fig. S1B,C**). In contrast, we noted a significant reduction in tail length, a surrogate for linear growth, in SD-IG relative to HD-IG mice (**Fig. 1C**). No difference in tail length was detected between SD-PW and HD-PW groups. Overall, both IG groups had significantly shorter tails than their PW counterparts, consistent with the finding of reduced body weight. As a second measure of linear growth, we also assessed femur length, which showed similar outcomes, although differences between SD-IG and HD-IG groups did not reach statistical significance (**Fig. 1D**). These trends were evident when comparing between IG and PW mice in each individual donor group and were not sex-dependent (**Fig. S1D-H**). Over the course of the experiment, we did not find significant differences in litter size between IG groups colonized with distinct donor microbiota (**Fig. S2A**). Because all four IG groups showed reduced growth compared to PW animals, these differences could reflect either paternal or early life M8 diet exposure in addition to earlier microbial colonization. These findings also suggest that intergenerational but not post-weaning colonization with SD microbiota negatively impacts linear growth relative to HD microbial communities.

### SD microbial communities influence intestinal histopathology

To begin to characterize underlying differences between SD-IG and HD-IG offspring that could explain the observed reductions in linear growth, we next examined small intestine histopathology. Blinded scoring of hematoxylin and eosin-stained ileal tissue sections from all four donor groups revealed significant reductions in villus length and muscularis thickness in SD-IG relative to HD-IG mice (**Fig. 1E**-**F**). Interestingly, these changes were not present when comparing HD-PW and SD-PW groups, suggesting that the reduction in villus length was not a general consequence of the presence of specific microbes, but depended on when these microbes were encountered. In addition to these quantitative measures, we also subjected sections to blinded scoring using three subjective parameters commonly observed in human intestinal biopsies from patients with EED ^16,30,31^, including immune cell infiltration, villous architecture and enterocyte injury. Surprisingly, we did not identify significant differences in these parameters, which may depend on additional environmental or pathogen exposures not replicated in our model (**Fig. S2B, Table S1A-B**).

### Distinct microbial communities colonize all four donor groups

In order to define the microbial communities mediating these effects, we next performed V4 16s rRNA sequencing of the fecal microbiota from mice in all four donor groups. Analysis of samples from IG animals at maturity revealed distinct community configurations for each donor (**Fig. 2A, Table S2A-D**), with a modest number of shared taxa (**Fig. S2C**). Analysis of UniFrac distances by nonmetric multidimensional scaling (NMDS) demonstrated that the composition of the fecal microbiota was similar between PW and IG samples within each individual donor, with the greatest variability present in donor 3114, and to a lesser extent, donor 4092 (**Fig. 2B, S2D**). Direct comparisons of PW to IG samples for each donor by weighted (**Fig. 2C**) and unweighted (**Fig. S2E**) UniFrac demonstrated significant differences in community structure between PW and IG mice colonized with the two stunted donor communities (**Table S2A-D**). Because PW mice were colonized directly while IG mice received vertically transmitted microbes, these results suggest the HD communities were efficiently passed from parents to offspring. In contrast, SD microbiota appeared to be surprisingly dependent on the mode of colonization (**Fig. 2D**). Whether these divergent patterns represent poor intergenerational transmission by the SD communities, more efficient engraftment by direct oral gavage, or shaping of the community by differential host immune responses remains to be determined.

**Figure 2.**
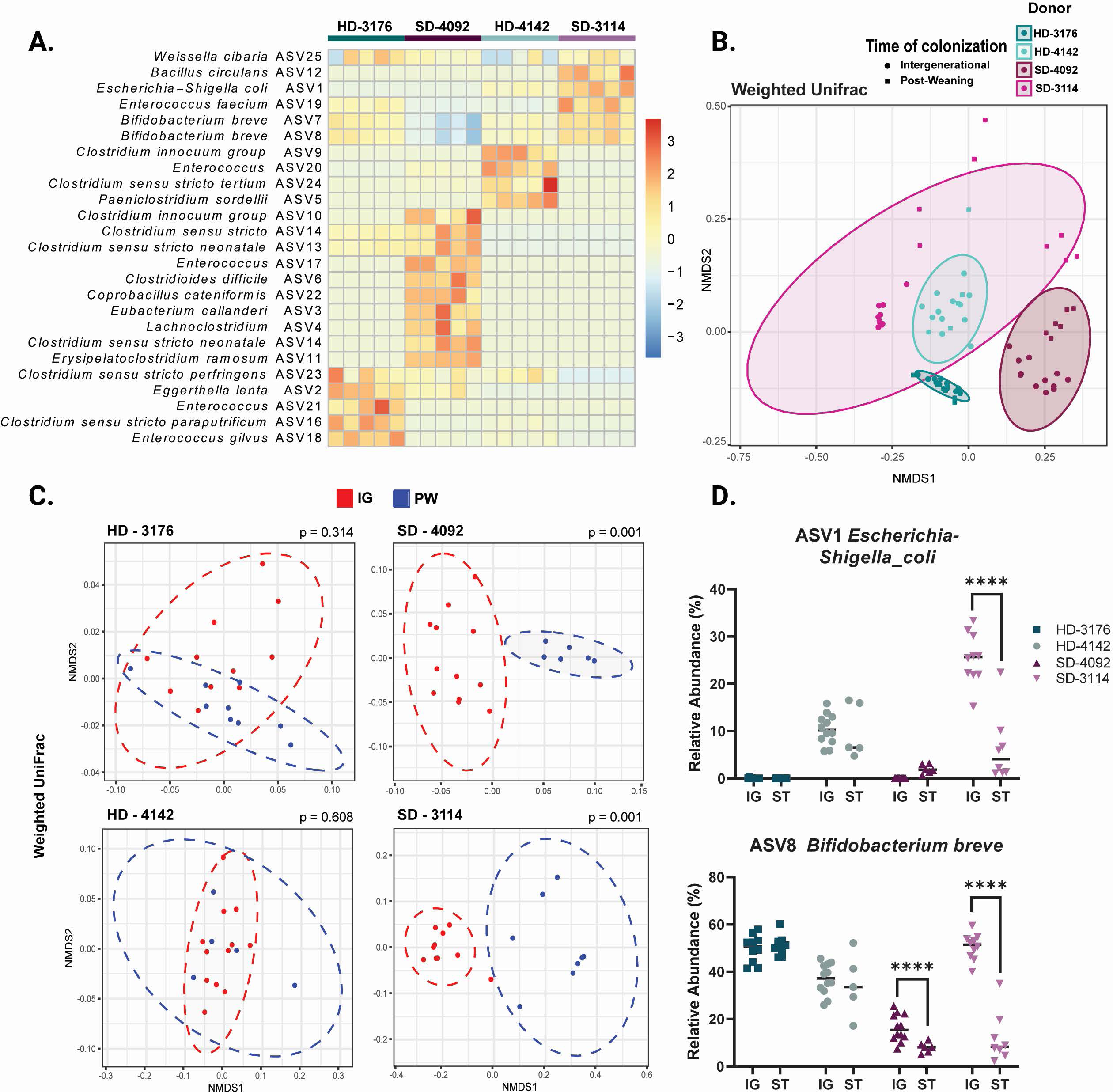
Microbial communities of recipient mice show distinct patterns of colonization. **(A)** Row-normalized heatmap of the top 25 ASVs by relative abundance for all four IG donor groups in fecal samples at 8 weeks of age. n=5/group. **(B)** NMDS plot of weighted UniFrac distances of IG and PW fecal samples at 8 weeks of age by donor colonization. n=5-12/group **(C)** NMDS plot of weighted UniFrac distances of IG versus PW fecal samples at 8 weeks of age by individual donor. n=5-11 mice/group with p value calculated by PERMANOVA. **(D)** Relative abundances of ASV1 *Escherichia-Shigella coli* and ASV 8 *Bifidobacterium breve* for all four IG and PW donor-colonized groups. n = 5-12/group, * p ≤ 0.05, ** p ≤ 0.01, **** p ≤ 0.0001 for comparisons between IG and PW samples within each donor by Mann-Whitney U test.

### Intergenerational colonization shapes small intestinal Intraepithelial Lymphocytes

Based on differences in linear growth and intestinal histopathology between SD-IG and HD-IG mice, we next sought to determine whether these microbial communities influenced immune cell composition within the small intestine epithelium. We first identified significantly elevated TCRβ+ cells in SD-IG relative to HD-IG mice (**Fig. 3A, S3A** and **Table S3A-D**), a difference that was not observed in HD-PW and SD-PW groups. Within this compartment, SD-IG mice had a significantly greater proportion of CD8α+ TCRβ+ cells and a significantly reduced proportion of CD4+ TCRβ+ T cells in the epithelium, whereas SD-PW and HD-PW mice did not (**Fig. 3B-C, S3B-C**). These results pointed towards potential differences in populations of intraepithelial lymphocytes (IELs), a heterogeneous group of microbiota-responsive immune cells located within the intestinal epithelium with diverse regulatory and inflammatory functions ^32–35^.

**Figure 3.**
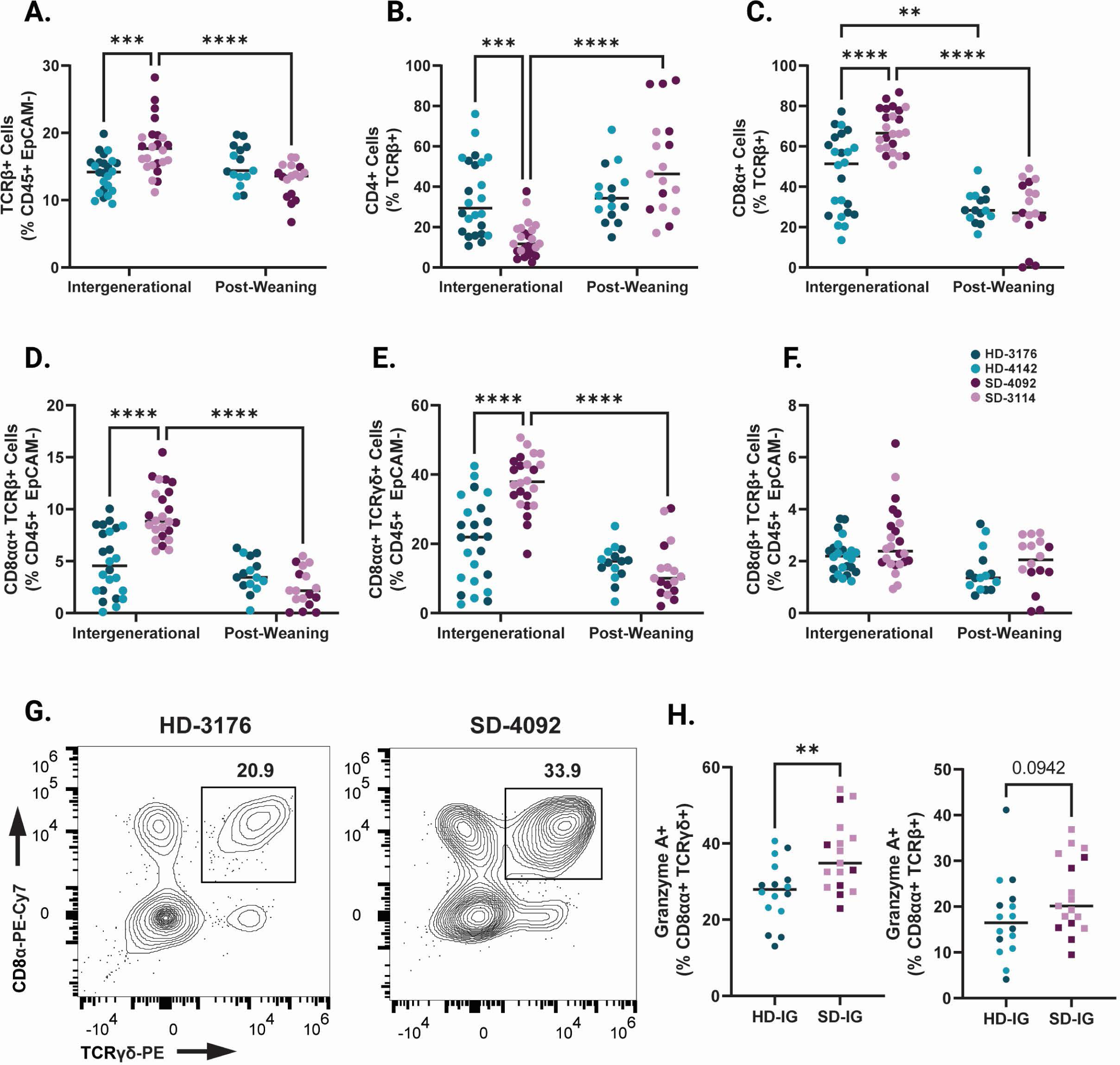
Immune cell composition of the small intestinal epithelium in IG and PW colonized mice at 8 weeks of age. **(A)** TCRβ+ cells shown as a percentage of CD45+ EpCAM-Live cells. **(B)** CD4+ cells shown as a percentage of TCRβ+ CD45+ EpCAM-Live cells. **(C)** CD8+ cells shown as a percentage of TCRβ+ CD45+ EpCAM-Live cells. **(D)** CD8αα+ TCRαβ Natural IELS (gated as CD8α+ CD8β-TCRβ+ cells as a percentage of CD45+ EpCAM-Live). **(E)** CD8αα+ TCRγδ Natural IELs (gated as CD8α+ CD8β-CD4-TCRγδ+ cells as a percentage of CD45+ EpCAM-Live). **(F)** CD8αβ+ TCRαβ Induced IELs (gated as CD8α+ CD8β+ TCRβ+ cells as a percentage of CD45+ EpCAM-Live). **(G)** Representative flow plots of TCRγδ+ Natural IELs in HD and SD IG mice. Cells were gated on the Live CD45+ EpCAM-population. **(H)** Granzyme A+ cells shown as a percentage of CD8α+ CD8β-CD4-TCRγδ+ CD45+ EpCAM-Live cells (left) or CD8α+ CD8β-TCRβ+ CD45+ EpCAM-Live cells (right) within IG colonized mice. Data shown pooled between healthy donor colonized mice (3176 and 4142) and stunted donor colonized mice (4092 and 3114). Each point represents an individual animal. (**A-F**) n = 24/group [12 per donor] for IG groups, n=18-20/group [7-10 per donor] for PW groups. * p ≤ 0.05, ** p ≤ 0.01, *** p ≤ 0.001, **** p ≤ 0.0001 by Two-Way ANOVA with Šídák’s multiple comparisons test. (**H**) n =16-17/group [6-11 per donor], ** p ≤ 0.01 by Mann-Whitney U test.

Findings from histological and molecular studies of human cohorts with EED have previously reported increases in IELs, and indeed, EED has been characterized as a T-cell-mediated enteropathy ^15,19,36,37^. IELs exist as two major subsets, including ‘natural’ IELs that are activated within the thymus, and ‘induced’ IELs that derive from conventional T cells activated peripherally ^33^. Intriguingly, SD-IG mice showed a significant increase in two subsets of natural IELs. CD8αα+ TCRβ+ and CD8αα+ TCRγδ+ IELs were both significantly more abundant in SD-IG mice relative to HD-IG animals, but neither cell type differed between the PW groups (**Fig. 3D-E, 3G, and Fig. S3D-E**). In contrast, induced CD8αβ+ TCRβ+ IELs did not significantly differ between groups (**Fig. 3F, S3F**). In keeping with recent results from single cell RNA sequencing of human intestinal biopsies during EED, a significantly greater proportion of CD8αα+ TCRγδ+ IELs were positive for Granzyme A, a cytotoxic mediator found in T lymphocytes (**Fig. 3H**) ^20^. Based on these results, we concluded that differences in natural IEL subsets were influenced by the composition of the microbiota when colonized intergenerationally, but not when colonized after weaning.

### Rorγt+ T regulatory cells are increased in SD-IG mice

We next investigated changes to innate and adaptive immune populations within the small intestine lamina propria. Overall, cellular changes in this compartment were less marked than those in the epithelium. However, we did note an overall increase in the proportions of Rorγt+ CD4+ T cells in the lamina propria of SD-IG mice (**Fig. 4A, Fig. S4A and Table S4A-D**). Recent findings from a murine model of EED induced by malnourished diet and adherent invasive *Escherichia coli* infection likewise reported increased microbiota-directed T regulatory cells, and increased numbers of these cells have also been reported in human cohorts with EED ^19,38^. Consistent with these findings, we found increased numbers of Rorγt+ FoxP3 regulatory T cells in the lamina propria of SD-IG mice relative to HD-IG animals (**Fig. 4B, S4B**). While the majority of innate immune cells were unchanged between groups, we did observe a reduction in lamina propria macrophages in SD-IG groups (**Fig. 4C, S4C**), again consistent with data from human cohorts with EED ^20^. Interestingly, the reduction in macrophages appeared to be driven by donor 4092 to a greater extent than donor 3114, suggesting differences in community composition between the two stunted donors may be involved. Finally, we also noted increased numbers of IgA+ Plasma Cells in the lamina propria of SD-IG mice relative to HD-IG mice (**Fig. 4D-E, S4D**). These cells were also significantly increased in HD-PW mice relative to HD-IG mice. Because the microbial communities of HD mice were similar between IG and PW groups, these results support the idea that the timing of exposure to the microbiota can play a major role in shaping the host immune response.

**Figure 4.**
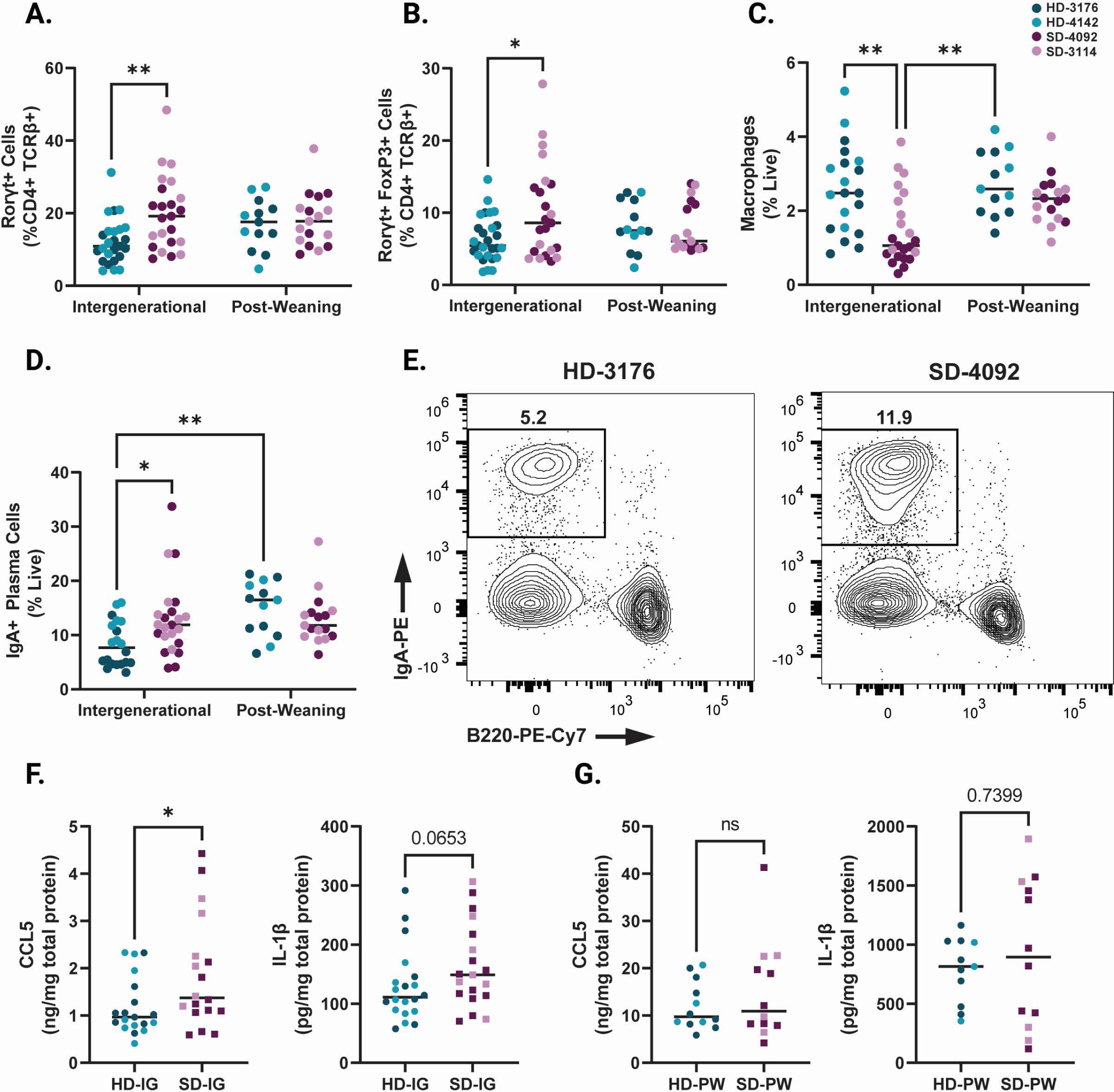
Immune features of the small intestine lamina propria in IG and PW colonized mice at 8 weeks of age. **(A)** Rorγt+ cells in the small intestine lamina propria shown as a percentage of CD4+ TCRβ+ live cells. **(B)** Rorγt+ Regulatory T cells (gated as Rorγt+ FoxP3+ CD4+ TCRβ+ live cells) in the small intestine lamina propria shown as a percentage of CD4+ TCRβ+ live cells. **(C)** Macrophages (gated as F4/80+ CD11b+ CD45+ live cells) in the small intestine lamina propria shown as a percentage of live cells. **(D)** IgA+ Plasma Cells (gated as Lin-[CD3-TCRβ-CD4-CD11c-NK1.1-F4/80-] IgA+ B220-live cells) in the small intestine lamina propria shown as a percentage of live cells. **(E)** Representative flow plots of IgA+ Plasma Cells (gated on live Lin-cells) in HD and SD IG groups. **(F)** Levels of CCL5 and IL-1β in ileal tissue lysates from IG mice as measured by ELISA. **(G)** Levels of CCL5 and IL-1β in ileal tissue lysates from PW mice as measured by ELISA. Data shown pooled between healthy donor colonized mice (3176 and 4142) and stunted donor colonized mice (4092 and 3114). Each point represents an individual animal. (**A-D**) n = 20-24/group [10-12 per donor] for IG groups, n=14-17/group [5-9 per donor] for PW groups. * p ≤ 0.05, ** p ≤ 0.01, *** p ≤ 0.001, **** p ≤ 0.0001 by Two-Way ANOVA with Šídák’s multiple comparisons test. (**F**) n =18-20/group [6-13 per donor], * p ≤ 0.05 by Mann-Whitney U test.

### SD-IG mice show elevated CCL5 and IL-1β in the small intestine

To investigate immune signals underlying the observed changes in cell composition in these groups, we next measured changes in small intestinal tissue chemokines and cytokines at the protein level. In ileum tissue lysates, we identified a significant increase in CCL5 (also known as regulated on activation, normal T cell expressed and secreted [RANTES]) protein by ELISA in SD-IG mice relative to HD-IG mice (**Fig. 4F, S4E-F**). CCL5 production can be induced by the microbiota in other murine models, notably exacerbating inflammation during DSS colitis ^39^. *CCL5* gene expression is also upregulated in duodenal biopsies obtained from a cohort of Pakistani patients with EED compared to healthy US controls or patients with celiac disease ^18^. Similarly, there was also a trend towards higher Interleukin-1β (IL-1β) in the SD-IG group. IL-1β is an acute phase protein whose secretion is triggered by activation of the inflammasome ^39,40^ (**Fig. 4G, S4G-H**). Overall, levels of these immune signaling molecules were markedly higher in both PW groups relative to both IG groups (**Fig. 4G-H**).

To further explore potential immune signaling differences between IG and PW colonization, we next performed multiplex bead-based analysis on a subset of ileal tissue lysate samples (**Table S5A-B**). Of the detectable cytokines and chemokines included in this analysis, we did not identify any additional significant differences between SD and HD groups in either the IG or PW samples. However, several were significantly elevated in both PW groups compared to both IG groups. These included IL-7, IL-10 and IL-17, among others (**Table S5A-B**). Overall, these results provide further insight into immune outcomes shaped by recognition of the microbiota in early life and support the idea that the timing of exposure to microbial communities plays a critical role in shaping immune outcomes.

## Discussion

Immune and epithelial changes in the small intestine are characteristic of EED and undernutrition, although the functional consequences of these changes are relatively poorly understood. Here we demonstrate that microbiota from two human infant donors with growth stunting can mediate reductions in linear growth and intestinal villus length as well as alterations in small intestinal immune cell populations relative to microbiota from healthy children. Differences between animals colonized with microbiota from these donors are evident when animals are born to colonized parents but are less apparent when animals are themselves directly colonized after weaning. These results are broadly consistent with previous work demonstrating that exposure of the immune system to microbial products during the weaning phase is a critical determinant of later life immune function ^27,28^. Our findings suggest that immune outcomes differ not only based on the presence of a microbiota during early life but also depends on which are microbes present.

These results raise questions about critical time points in early life that shape growth and immunity. While the current body of evidence suggests the weaning phase is one critical time period, many children who develop stunting show reduced linear growth at birth, and our results do not rule out a potential role for the maternal microbiome during development *in utero* ^11^. Furthermore, a weaning reaction to the microbiota has not been demonstrated in humans, so it is unclear how these findings translate to children with undernutrition and EED. Despite these challenges, we identified certain immune alterations in our IG model (increased IELs, regulatory T cells and plasma cells) that have also been shown in patients with EED ^18–20^, suggesting certain features of the immune response to the microbiota are shared between mice and humans. In summary, we suggest this model may be a useful system with which to elucidate the functional role of immune alterations during undernutrition and EED in early life.

## Limitations of our study

There are several important caveats to the current study. First, due to the complexity of the experimental design employed, we were unable to investigate more than four total microbiota donors. While several of our findings show similarities to human studies in patients with undernutrition and EED, the low number of donors employed limits the overall generalizability of our results. We likewise limited our investigation to microbiota samples from one specific age group (six-month-old infants), and it is unclear how later diversification of the microbiome in children with varied environmental exposures could impact immune composition and function at maturity. Another challenge in interpreting our results is the finding that colonization of the two SD microbiota groups differed significantly between PW and IG mice. While interesting, this finding does suggest that observed differences in PW and IG SD mice could be driven by differences in community composition rather than timing of colonization. It also remains to be determined whether this model of undernutrition can recapitulate other long-term sequelae of undernutrition, including oral vaccine failure, increased susceptibility to infectious disease, and cognitive developmental changes. These possibilities warrant further investigation.

## Acknowledgements

We thank UVA’s Gnotobiotic and Germ-Free Animal Facility and Rory Laman for their help in performing these experiments, which would not have been possible without their dedicated efforts. We thank the participants and investigators of the iLiNS-Dyad-M study for the samples used in this study (provided under material transfer agreement from Washington University in St. Louis to the University of Virginia). We thank UVA’s Flow Cytometry Core Facility (RRID: SCR_017829) and Research Histology Core for their contributions to this work. Y.S.M. was supported by NIH T32 AI055432 and NSF LSAMP Bridge to the Doctorate Fellowship 1810762. J.C. and C.A.C. were supported by NIH R01 HD105729. C.W. was supported by NIH T32 AI007496.

## Author Contributions

Conceptualization, Y.S.M and C.A.C.; Sample collection, Y.S.M, J.C., H.S., C.W., J.S. and C.A.C.; Investigation, Y.S.M, J.C., H.S., C.W., J.S. and C.A.C., Analysis; Y.S.M, J.C., H.S., J.S. and C.A.C., Visualization, Y.S.M, J.C., H.S., J.S. and C.A.C.; Resources, J.C., Y.S.M and C.A.C; Funding, Y.S.M. and C.A.C, Supervision, C.A.C; Writing, Y.S.M, J.C. and C.A.C.

## Figure Titles/Legends

**Supplemental Figure 1.**
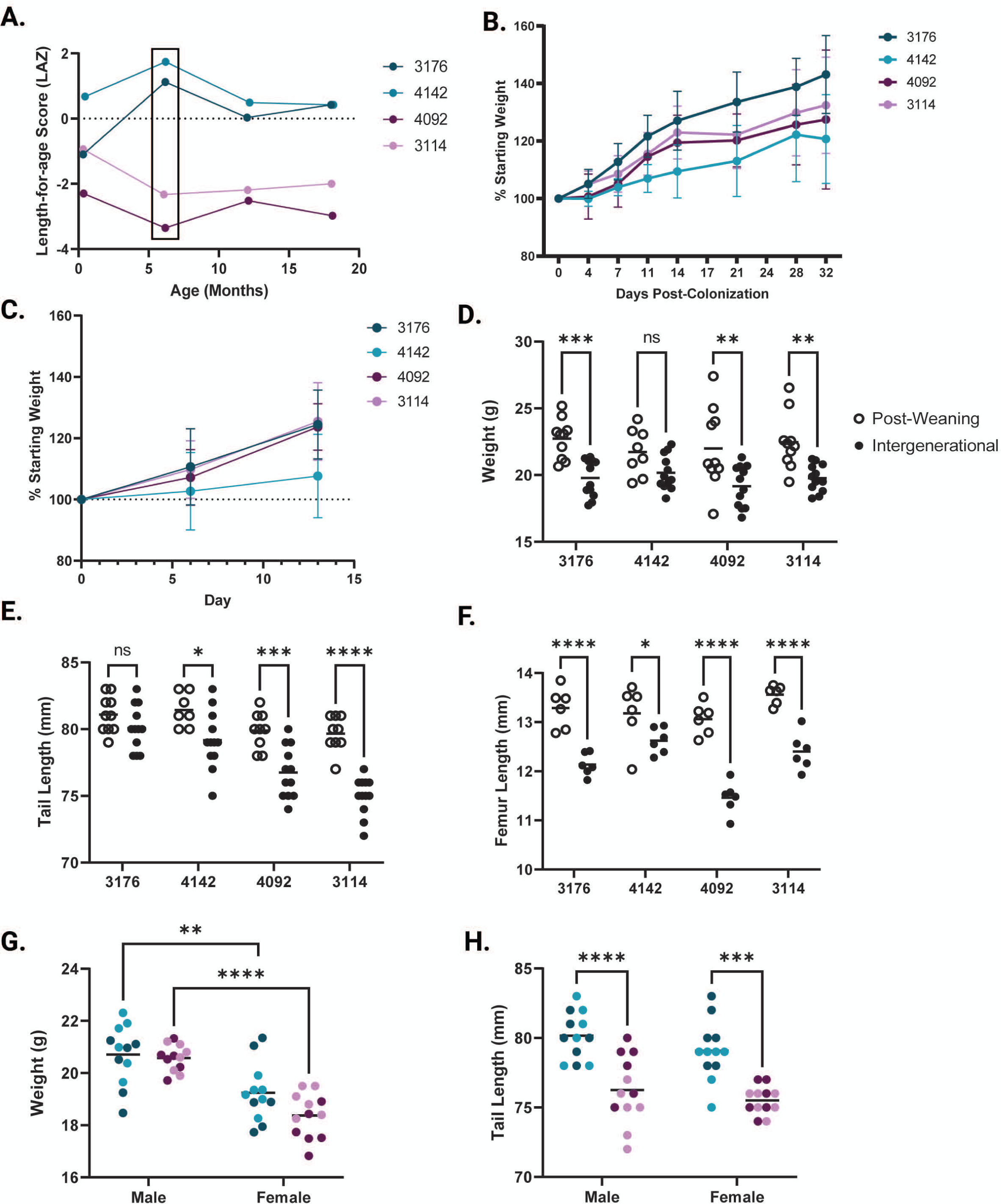
Donor and recipient growth parameters. **(A)** Length-for-age Z scores from microbiota donors over the first 20 months of life ^23,29^. Microbiota samples used in this study were collected at the six-month time point. **(B)** Weight of recipient gnotobiotic mice colonized with individual microbiota donors at four weeks of age (post-weaning mice), shown as a percentage of starting weight. n=6/group, evaluated statistically by Mixed-effects model with Tukey’s multiple comparisons test. **(C)** Weight of recipient gnotobiotic mice colonized with individual microbiota donors at four weeks of age (sires of intergenerational mice), shown as a percentage of starting weight. n=5-7/group, evaluated statistically by Mixed-effects model with Tukey’s multiple comparisons test. **(D)** Absolute weight of PW and IG mice by individual microbiota donor at 8 weeks of age. **(E)** Tail length measurements of PW and IG mice by individual microbiota donor at 8 weeks of age. n = 12/group for IG groups, n=8-10/group for PW groups in **D-E**. **(F)** Femur length measurements of PW and IG mice by individual microbiota donor at 8 weeks of age. n=6/group. **(G)** Comparison of male and female HD and SD IG weights at 8 weeks of age. **(H)** Comparison of male and female tail lengths for HD and SD IG mice at 8 weeks of age. n=12/group [6 per donor], ** p ≤ 0.01, *** p ≤ 0.001, **** p ≤ 0.0001 by Two-Way ANOVA with Šídák’s multiple comparisons test for **G** and **H**.

**Supplemental Figure 2.**
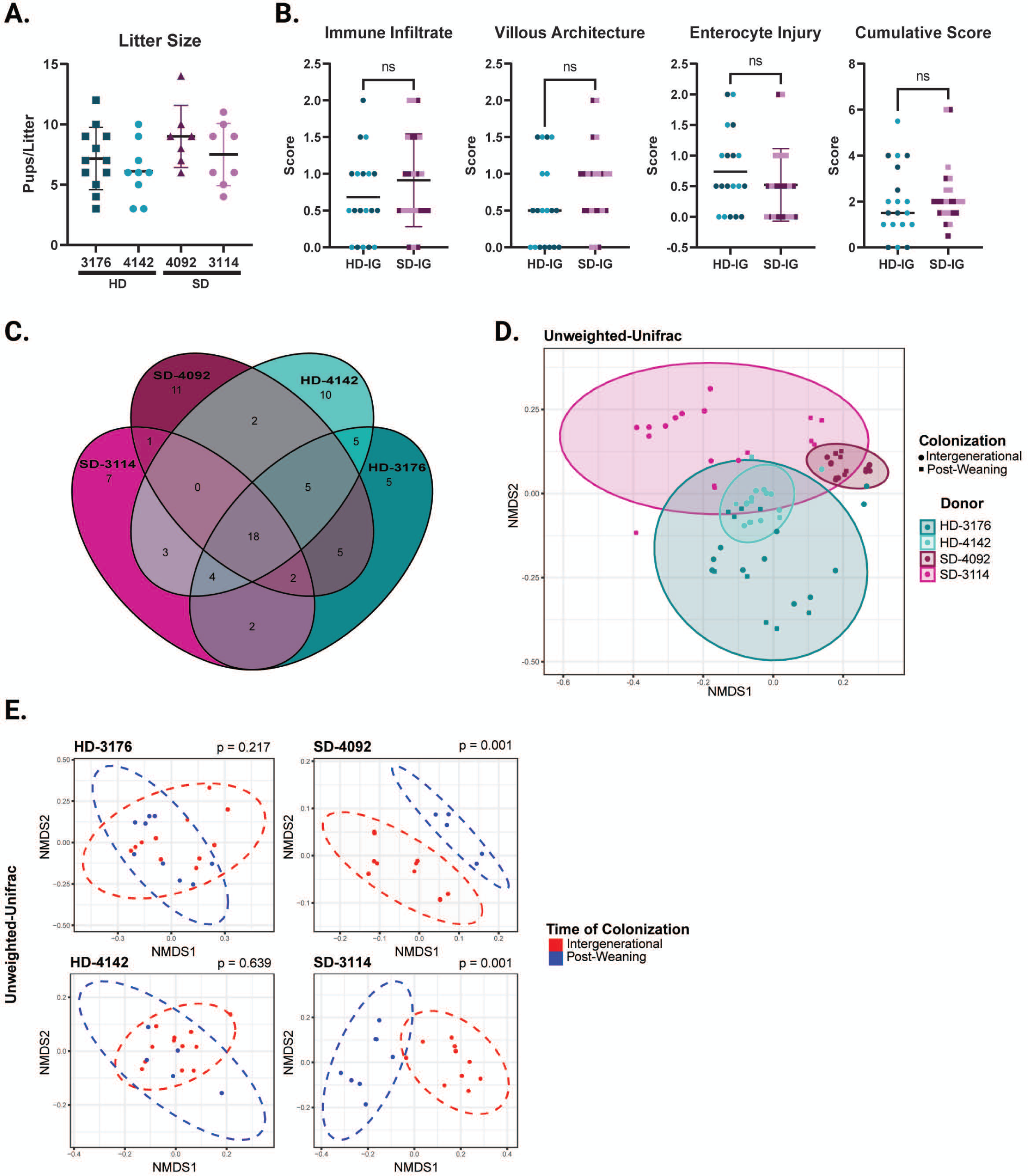
Breeding statistics, histological parameters, and microbiota composition of HD and SD IG mice. **(A)** Litter size for all litters born to IG breeding pairs over the course of the experiment. No significant differences were identified by Kruskal-Wallis test with Dunn’s multiple comparisons. **(B)** Subjective histological scoring parameters for HD and SD IG mice, shown averaged between two independent blinded observers. n =18-22/group [7-12 per donor] for IG groups, evaluated by Mann-Whitney U test. **(C)** Venn diagram of unique and shared ASVs identified by V4 16s sequencing of fecal microbiota from all four IG groups at 8 weeks of age. n=10-12 per group. **(D)** NMDS plot of unweighted UniFrac distances of IG and PW fecal samples at 8 weeks of age by donor colonization. n=5-12/group **(E)** NMDS plot of unweighted UniFrac distances of IG versus PW fecal samples at 8 weeks of age by individual donor with p value calculated by PERMANOVA. n=5-11 mice/group

**Supplemental Figure 3.**
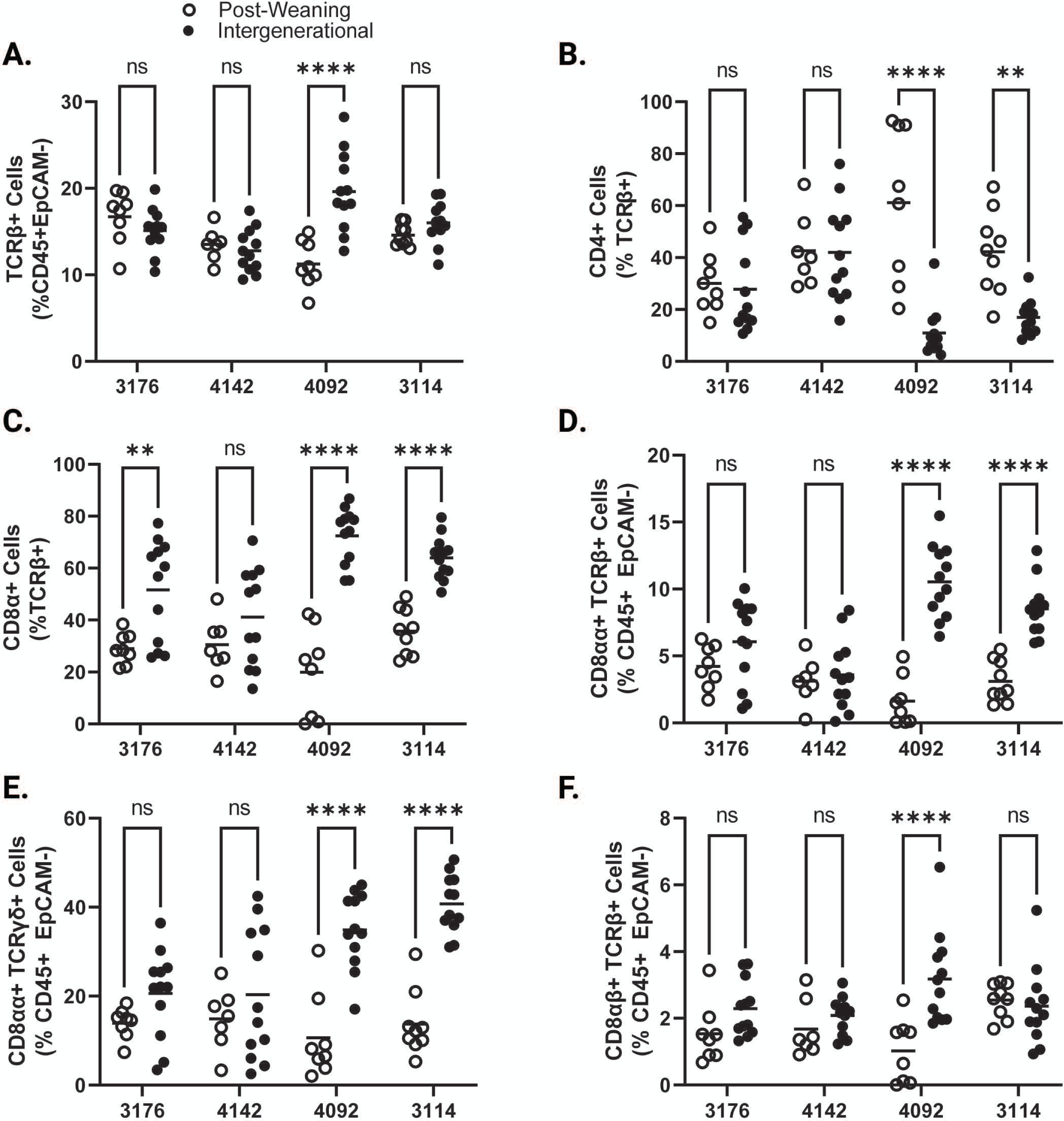
Immune cell composition of the small intestine epithelium at 8 weeks of age for each individual microbiota donor by timing of colonization. **(A)** TCRβ+ cells shown as a percentage of CD45+ EpCAM-live cells. **(B)** CD4+ cells shown as a percentage of TCRβ+ CD45+ EpCAM-live cells. **(C)** CD8+ cells shown as a percentage of TCRβ+ CD45+ EpCAM-live cells. **(D)** CD8αα+ TCRαβ Natural IELS (gated as CD8α+ CD8β-TCRβ+ cells as a percentage of CD45+ EpCAM-Live). **(E)** CD8αα+ TCRγδ Natural IELs (gated as CD8α+ CD8β-CD4-TCRγδ+ cells as a percentage of CD45+ EpCAM-Live). **(F)** CD8αβ+ TCRαβ Induced IELs (gated as CD8α+ CD8β+ TCRβ+ cells as a percentage of CD45+ EpCAM-Live). (**A-F**) Each point represents an individual animal. n = 7-12/group * p ≤ 0.05, ** p ≤ 0.01, *** p ≤ 0.001, **** p ≤ 0.0001 by Two-Way ANOVA with Šídák’s multiple comparisons test.

**Supplemental Figure 4.**
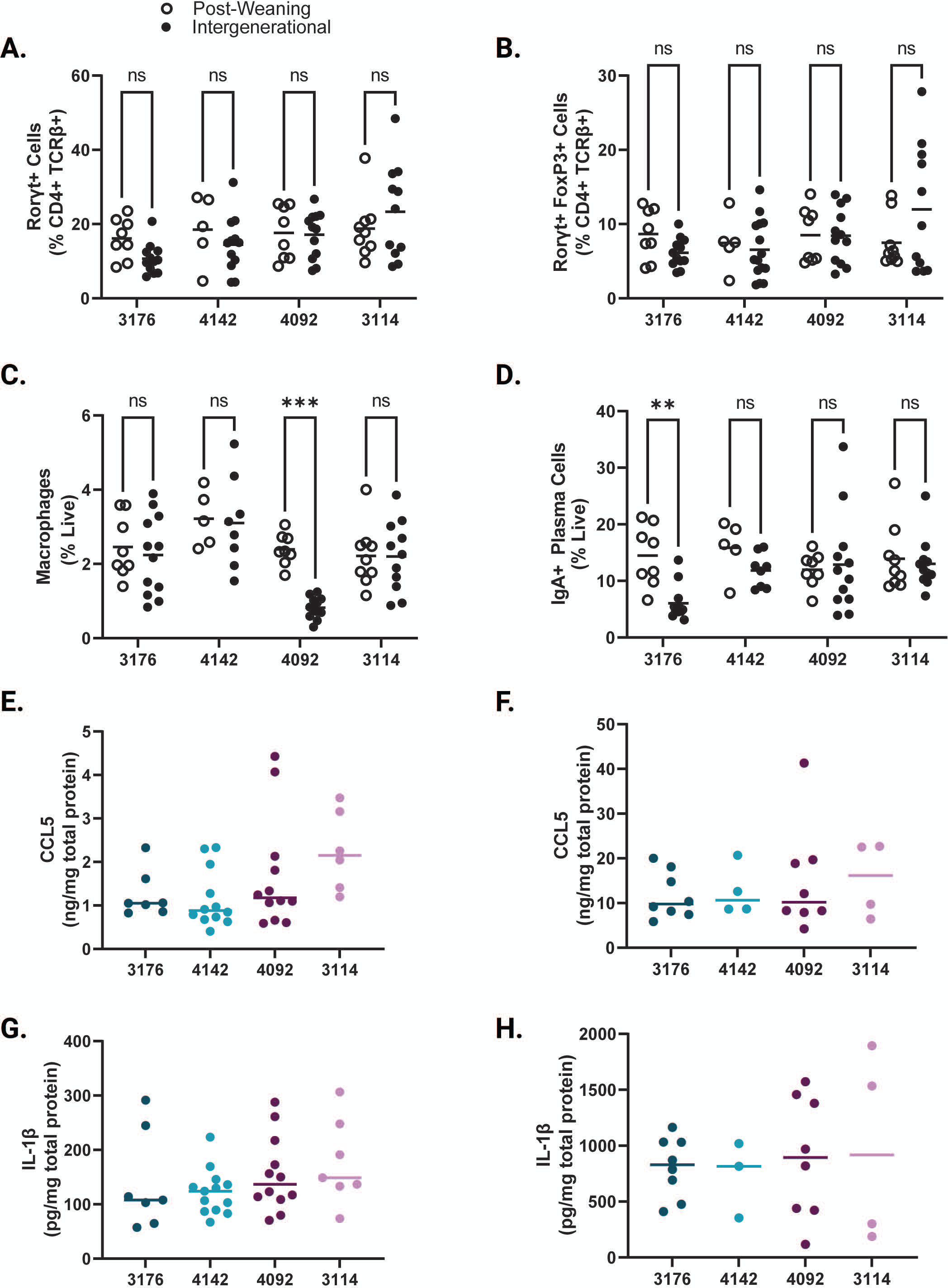
Immune features of the small intestine lamina propria in IG and PW colonized mice at 8 weeks of age for each individual microbiota donor by timing of colonization. **(A)** Rorγt+ cells in the small intestine lamina propria shown as a percentage of CD4+ TCRβ+ live cells. **(B)** Rorγt+ Regulatory T cells (gated as Rorγt+ FoxP3+ CD4+ TCRβ+ live cells) in the small intestine lamina propria shown as a percentage of CD4+ TCRβ+ live cells. **(C)** Macrophages (gated as F4/80+ CD11b+ CD45+ live cells) in the small intestine lamina propria shown as a percentage of live cells. **(D)** IgA+ Plasma Cells (gated as Lin-[CD3-TCRβ-CD4-CD11c-NK1.1-F4/80-] IgA+ B220-live cells) in the small intestine lamina propria shown as a percentage of live cells. **(E)** Levels of CCL5 in ileal tissue lysates from IG mice as measured by ELISA. n =7-13/group **(F)** Levels of IL-1β in ileal tissue lysates from IG mice as measured by ELISA. n =7-13/group **(G)** Levels of CCL5 in ileal tissue lysates from PW mice as measured by ELISA. n =4-8/group **(H)** Levels of IL-1β in ileal tissue lysates from PW mice as measured by ELISA. n =3-8/group **(A-D)** Each point represents an individual animal. n = 12/group for IG groups, n=5-9/group for PW groups. * p ≤ 0.05, ** p ≤ 0.01, *** p ≤ 0.001, **** p ≤ 0.0001 by Two-Way ANOVA with Šídák’s multiple comparisons test.

**Supplemental Table 1. Histological scoring parameters of H&E stained ileal tissue from 8 week old mice at the time of euthanasia.**

**(A)** Histological assessment of H&E stained ileal tissue harvested from gnotobiotic mice at the time of euthanasia, expressed by microbiota donor health status.

**(B)** Histological assessment of H&E stained ileal tissue harvested from gnotobiotic mice at the time of euthanasia, expressed by individual microbiota donor.

**(A-B)** n=8-12/group (IG samples) or 8-10/group (PW samples). * p ≤ 0.05, ** p ≤ 0.01, *** p ≤ 0.001, **** p ≤ 0.0001 by Mann-Whitney U test

**Supplemental Table 2. Relative abundances of the 25 most abundant ASVs per donor by V4 16s Sequencing of fecal samples at 8 weeks of age.**

**(A)** Top 25 most abundant microbial taxa across intergenerational and post-weaning mice colonized with donor microbiota 3176, expressed as percent relative abundance.

**(B)** Top 25 most abundant microbial taxa across intergenerational and post-weaning mice colonized with donor microbiota 4142, expressed as percent relative abundance.

**(C)** Top 25 most abundant microbial taxa across intergenerational and post-weaning mice colonized with donor microbiota 4092, expressed as percent relative abundance.

**(D)** Top 25 most abundant microbial taxa across intergenerational and post-weaning mice colonized with donor microbiota 3114, expressed as percent relative abundance.

**(A-D)** n=10-12/group (IG samples) or 5-9/group (PW samples). * p ≤ 0.05, ** p ≤ 0.01, *** p ≤ 0.001, **** p ≤ 0.0001 by Mann-Whitney U test

**Supplemental Table 3. Immune populations in the small intestine epithelium of gnotobiotic mice colonized with healthy or stunted donor microbial communities.**

**(A)** Immune cell composition of the small intestine epithelium harvested from gnotobiotic mice at the time of euthanasia, expressed as percentages by microbiota donor health status.

**(B)** Immune cell composition of the small intestine epithelium harvested from gnotobiotic mice at the time of euthanasia, expressed as total number of cells per intestine.

**(C)** Immune cell composition of the small intestine epithelium harvested from gnotobiotic mice at the time of euthanasia, expressed as percentages by individual microbiota donor.

**(D)** Immune cell composition of the small intestine epithelium harvested from gnotobiotic mice at the time of euthanasia, expressed as total number of cells per intestine by individual microbiota donor.

Each point represents an individual animal. (**A-D**) n = 24/group [12 per donor] for IG groups, n=18-20/group [7-10 per donor] for PW groups. * p ≤ 0.05, ** p ≤ 0.01, *** p ≤ 0.001, **** p ≤ 0.0001 by Two-Way ANOVA with Šídák’s multiple comparisons test.

**Supplemental Table 4. Immune populations in the small intestine lamina propria of gnotobiotic mice colonized with healthy or stunted donor microbial communities.**

**(A)** Immune cell composition of the small intestine lamina propria harvested from gnotobiotic mice at the time of euthanasia, expressed as percentages by microbiota donor health status.

**(B)** Immune cell composition of the small intestine lamina propria harvested from gnotobiotic mice at the time of euthanasia, expressed as total number of cells per intestine by microbiota donor health status.

**(C)** Immune cell composition of the small intestine lamina propria harvested from gnotobiotic mice at the time of euthanasia, expressed as percentages by individual microbiota donor.

**(D)** Immune cell composition of the small intestine lamina propria harvested from gnotobiotic mice at the time of euthanasia, expressed as total number of cells per intestine by individual microbiota donor.

Each point represents an individual animal. (**A-D**) n = 24/group [12 per donor] for IG groups, n=13-17/group [5-9 per donor] for PW groups. * p ≤ 0.05, ** p ≤ 0.01, *** p ≤ 0.001, **** p ≤ 0.0001 by Two-Way ANOVA with Šídák’s multiple comparisons test.

**Supplemental Table 5. Cytokine and chemokine levels in the ileum of gnotobiotic mice colonized with healthy or stunted donor microbial communities.**

**(A)** Multiplex bead analysis of ileal tissue lysate from gnotobiotic mice at the time of euthanasia, expressed as pg/mg total protein by microbiota donor health status.

**(B)** Multiplex bead analysis of ileal tissue lysate from gnotobiotic mice at the time of euthanasia, expressed as pg/mg total protein by individual microbiota donor.

Each point represents an individual animal. (**A-D**) n = 5-6/group, * p ≤ 0.05, ** p ≤ 0.01, *** p ≤ 0.001, **** p ≤ 0.0001 by Two-Way ANOVA with Šídák’s multiple comparisons test.

## Methods

### Lead contact

Further information and requests for resources and reagents should be directed to and will be fulfilled by the lead contact, Carrie Cowardin.

### Materials availability

This study did not generate new unique reagents.

### Data and code availability

• V4 16s rRNA sequencing data will be deposited in the European Nucleotide Archive prior to publication.

• This paper does not report original code. Analysis tools for 16s rRNA sequencing data are listed in the key resources table and are publicly available.

• Any additional information required to reanalyze the data reported in this paper is available from the lead contact upon request.

## EXPERIMENTAL MODEL DETAILS

### Donor microbiota and study protocol

Details of enrollment for the iLiNS-DYAD-M study were described in an earlier publication ^23^. Briefly, enrollment was open to consenting women over the age of 15 years with ultrasound confirmation of pregnancy of <20 weeks gestation in the Mangochi District of southern Malawi. The randomized controlled clinical trial [clinicaltrials.gov #NCT01239693] tested the effects of providing small quantity Lipid-based Nutrient Supplements (SQ-LNS) to pregnant and lactating women through 6 months postpartum and to their children through 6-18 months of age ^29^. Donors used in this study were selected from a broader subset of donors based on their ability to colonize recipient gnotobiotic mice at an efficiency of >50% (more than half of taxa present in the original donor sample were identified in initial experiments ^23^).

### Gnotobiotic Mice

All mouse experiments were performed using protocols approved by the University of Virginia Institutional Animal Care and Use Committee. All animals used in this publication were germ-free C57Bl6 mice obtained from Taconic Biosciences. Upon arrival, germ-free status was verified by quantitative PCR. Mice were housed in plastic flexible film gnotobiotic isolators (Class Biologically Clean Ltd., Madison WI) under a 12-hour light cycle. Animals received *ad libitum* access to food and water throughout the experiment, and were euthanized at the conclusion of the experiment using AVMA approved procedures. For Post-Weaning experiments, male and female mice were obtained from Taconic Biosciences at 4 weeks of age, and were immediately transitioned to the M8 diet upon arrival. Three days later, they were colonized with donor microbiota and maintained thereafter on M8 until the time of euthanasia at 8 weeks of age. For Intergenerational experiments, male mice were obtained from Taconic Biosciences at 4 weeks of age and were immediately transitioned to the M8 diet upon arrival. Three days later, they were colonized with donor microbiota and maintained on M8 diet until 8 weeks of age. At this point, 8 week old germ-free females were introduced into the donor isolators. Males and females were co-housed and switched to an autoclaved nutrient-sufficient breeder chow (LabDiet 5021 Autoclavable Mouse Breeder Diet, LabDiet Inc.). Breeding animals were maintained on this diet thereafter. Offspring of these breeders were weaned at 21 days of life onto the M8 diet, and maintained on this diet until euthanasia at 8 weeks of age. Breeding mice were refreshed after six months, and offspring used in these experiments were derived from breeders from two separate rounds of colonization (total of 4 males and 6 females per breeding isolator). All data shown represent results from a minimum of two separate litters.

### Colonization and Diets

To prepare infant fecal samples for colonization of germ-free mice, aliquots of each sample were removed from storage at −80°C, weighed and immediately transferred into anaerobic conditions (atmosphere of 75% N2, 20% CO2 and 5% H2; vinyl anaerobic chambers from Coy Laboratory Products). Samples were subsequently resuspended in pre-reduced PBS containing 0.05% L-Cysteine Hydrochloride at a concentration of 10 mg/mL. Samples were vortexed for one minute and allowed to clarify by gravity for 5 minutes. The supernatant was removed to a fresh anaerobic tube and combined with an equal volume of sterile, pre-reduced PBS containing 0.05% L-Cysteine HCL and 30% glycerol. Gavage mixtures were aliquoted into sterile 2 mL screw cap tubes (Axygen) and frozen at −80°C until use. To colonize recipient mice, pools of gavage mixtures were sterilized externally with ionized hydrogen peroxide (STERAMIST System, TOMI Inc.) and passed into each isolator after appropriate exposure time (20 minutes). Animals were colonized via a single oral gavage with a 200 µl volume of gavage mixture.

The micro and macro-nutrient deficient Malawi-8 diet was prepared as previously described ^23,41^ and obtained from Dyets, Inc. Briefly, ingredients (corn flour, mustard greens, onions, tomatoes, ground peanuts, red kidney beans, canned pumpkin and peeled bananas) were cooked and combined in an industrial mixer. Dry pellets of the M8 diet were extruded, vacuum-sealed and double bagged prior to sterilization by irradiation (Steris Co; Chicago, IL). The nutritional content of the cooked and irradiated diet was assessed by N.P. Analytical Laboratories (St. Louis, MO) as described in Blanton et al ^23^. LabDiet 5021 was sterilized by autoclaving at 129°C and 13.2 PSI for 15 minutes. Sterility of both diets was routinely assessed by culturing pellets in Brain Heart Infusion (BHI) broth, Nutrient broth, and Sabouraud-Dextrose broth (Difco) for five days at 37 °C under aerobic conditions, and in BHI broth and Thioglycolate broth (Difco) supplemented with 0.05% L-Cysteine Hydrochloride (Sigma) under anaerobic conditions. After the five-day liquid culture, cultures of all diets were plated on BHI agar supplemented with sheep blood (Thermo Scientific). All diets were stored at −20 °C prior to use.

## METHOD DETAILS

### Histology

At the time of euthanasia, a 1 cm section of the proximal ileum was dissected from each mouse and fixed in 10% neutral-buffered formalin overnight at room temperature before being transferred to 70% Ethanol. Tissue processing and H&E staining were performed by the University of Virginia’s Research Histology Core. Samples were paraffin embedded and sectioned before mounting. Slides were stained with hematoxylin & eosin prior to imaging at a 20x magnification using an EVOS M7000 microscope. To assess the histopathological features of the ileum, the stained tissues were scored in a blinded manner by two independent observers. Resulting scores were then averaged. Scores were assigned using a scoring system based off published findings in human intestinal biopsies ^30^. Scoring parameters consisted of three qualitative features: immune cell infiltration (0, no visible increase in tissue area; 1, increase in immune cells present in < 50% of tissue area; 2, increase in immune cells present in > 50% of tissue area), villous architecture (0, majority of villi are >3 crypt lengths long; 1, majority of villi are <3 but >1 crypt length long, with abnormality; 2, majority of villi are absent or <1 crypt length long, with abnormality) and enterocyte injury (0, majority of enterocytes show tall columnar morphology; 1, < 50% of enterocytes show low columnar, cuboidal, or flat morphology; 2, > 50% of enterocytes show low columnar, cuboidal, or flat morphology). Cumulative scores were calculated as the sum of the averaged score for all three parameters. ImageJ was used to obtain two quantitative parameters consisting of ileum villus height (μm) and ileum muscularis thickness (μm). Two measurements were obtained for each parameter and averaged.

### Microbial Sequencing

DNA was prepared from fecal samples by bead beating (BioSpec Products) in a solution containing 500µL of extraction buffer [200 mM Tris (pH 8.0), 200 mM NaCl, 20 mM EDTA], 210µL of 20% SDS, 500µL phenol:chloroform:isoamyl alcohol (pH 7.9, 25:24:1, Calbiochem), and 500µL of 0.1-mm diameter zirconia/silica beads. The aqueous phase was removed and DNA purified by PCR Purification Kit (Qiagen). Pure DNA was quantified by Qubit dsDNA BR assay (Invitrogen). DNA was normalized to a 2 ng/ul concentration and 15 ng total DNA was used as template for subsequent PCR reactions. Bacterial V4 16S rRNA gene amplicons were generated using custom barcoded primers ^42^. PCR was performed with Invitrogen High Fidelity Platinum Taq using the manufacturer’s suggested cycling conditions. No-template controls were run with every sample plate to ensure there was no contamination of the barcoded primers or reagents. Amplicons were purified using Qiagen Qiaquick Purification Kit following the manufacturer’s protocol. To confirm the presence of targeted amplicons, PCR products were subjected to gel electrophoresis followed by quantification using the Qubit hsDNA Assay (Invitrogen). Barcoded amplicons were pooled to a concentration of 4 nM then sequenced using the Miseq Reagent Kit v3 and Miseq platform (Illumina) following the manufacturer’s recommended protocols.

### Sequencing Quality Control & Data Analysis

Sequence reads were demultiplexed using Illumina Miseq Reporter Software. Quality checks were performed using fastqc version 0.11.5 before and after trimming. Samples with less than total 5,000 reads were excluded from downstream analysis. Remaining reads were trimmed using the Bbduk tool from BBMap (https://sourceforge.net/projects/bbmap/) to 150 base pairs (bp) and with sequence quality qc ≥10. ASVs were produced using customized DADA2 pipeline. Taxonomic assignment was performed using the SILVA ribosomal RNA gene database. ASVs were filtered to exclude taxa with relative abundance across all samples < 0.1% or prevalence across samples < 2%. Ordination plots and heat maps were produced using the phyloseq package in R ^43^.

### Flow Cytometry

#### Tissue Harvest and Cell Isolation

The entire small intestine was dissected from each mouse and placed onto a moist piece of gauze soaked in HBSS with 10 mM HEPES (Gibco) to prevent drying. The intestine was separated into duodenum, jejunum, and ileum, and a small portion (∼1 cm) of the proximal end of each section was removed for histological analysis. Intestinal contents were then gently squeezed out of each section. Peyer’s Patches and any fat were removed from the small intestine exterior, then the tissue was sliced lengthwise to expose the lumen, and samples were placed in 10ml of HBSS with 10 mM HEPES on ice. Intestines were then transferred to 20ml of room temperature epithelial removal buffer (1x HBSS with 20% FBS, 7.5 mM HEPES, and 2.5 mM EDTA) and rotated on a tube rotator for 20 minutes at room temperature, then vortexed for 20 seconds. Intestines and buffer were then poured over a 100 µm filter and the collected cell suspension containing intestinal epithelial cells was spun at 500xg for 5 minutes at 4°C, resuspended in 5ml FACS buffer (1x DPBS with 2% FBS), and reserved for flow cytometric staining and analysis. This process was repeated once more with each intestine sample and the remaining intact tissue at this point consisted of small intestine lamina propria. Tissue was then rinsed in 1x HBSS with 10 mM HEPES to remove residual epithelial removal buffer, and finely chopped with scissors, then resuspended in 10ml of warmed complete RPMI (Gibco) and 100ul of 100 U/mL collagenase IV (Millipore Sigma). Samples were incubated at 37°C on a shaker at 180rpm for 45 minutes. The resulting mixture was poured over a 40 µm filter and tissue mashed through the filter with the end of a syringe plunger. Plunger, filter, and original digestion tube were rinsed with up to 5ml of FACS buffer to ensure maximum cell yield. Resulting cell suspension was spun at 500xg for 5 minutes at 4°C, then the supernatant was decanted, and the cell pellet was resuspended in 4mls of 40% Percoll (Cytiva) in HBSS in a 15ml conical tube. This Percoll layer was gently underlaid with 3ml of 70% Percoll in HBSS to create two distinct layers. Tubes were spun at 850xg for 20 minutes at 4°C with the centrifuge brake turned off to prevent mixing of the Percoll layers. After spin, the interface of cells between the two Percoll layers was collected and added to 10ml of HBSS with 10 mM HEPES. Tubes were spun at 500xg for 5 minutes at 4°C, then supernatant was aspirated off and cell pellets were resuspended in 500ul of FACS buffer.

#### Surface and Intracellular Staining for Flow Cytometry

Epithelium and lamina propria cell suspensions were plated in 96 well round bottom plates, spun at 500xg at 4°C for 5 minutes, supernatant decanted, and 50ul of surface stain mixture (fluorescent antibodies for surface markers, TruStain FcX PLUS [Biolegend], and Zombie Aqua Fixable Viability dye [Biolegend]) in FACS buffer was applied to each well and incubated for 20 minutes at room temperature in the dark. Plates were then spun at 500xg at 4°C for 5 minutes, and supernatant decanted. If cells were receiving no intracellular stains, cells were resuspended in 100ul of 1x Fixation Buffer (BD Biosciences) for 20 minutes in the dark at 4°C. They were then spun at 500xg at 4°C for 5 minutes, supernatant was decanted, and they were resuspended in 200ul of FACS buffer and allowed to sit at 4°C overnight in the dark. On the day of analysis, cells were spun at 500xg at 4°C for 5 minutes, supernatant decanted, and resuspended in 200ul FACS buffer plus 5ul of CountBright Absolute Counting Beads (Thermo Fisher). For intracellular staining, cells were fixed in 100ul FoxP3 Fix/Perm Buffer (Invitrogen) for 20 minutes at 4°C in the dark, then spun at 500xg at 4°C for 5 minutes, supernatant was decanted, and cells were resuspended in 100ul of intracellular antibodies diluted in 1x permeabilization buffer (Invitrogen) with 2% rat serum (Invitrogen) and incubated overnight at 4°C in the dark. On the day of analysis, cells were spun at 500xg at 4°C for 5 minutes, supernatant decanted, resuspended in 200ul of 1x permeabilization buffer and allowed to sit at room temperature for 5 minutes. Cells were then spun again, decanted, and resuspended in 200ul FACS buffer plus 5ul of CountBright Absolute Counting Beads (Thermo Fisher).

#### Analysis and Gating

Samples were run on the Attune NxT Acoustic Focusing Cytometer with CytKick Auto Sampler and analyzed using FlowJo software. All cell types were first gated as live, single cells. Subsequent gating was performed using the following parameters and antibodies:

| Tissue Type | Cell Type | Markers |
| --- | --- | --- |
| Epithelium | $\alpha\beta$ T Cells | CD45+ EpCAM- TCRβ+ |
| Epithelium | CD4+ T Cells | CD45+ EpCAM- TCRβ+ CD4+ |
| Epithelium | CD8+ T cells | CD45+ EpCAM- TCRβ+ CD8+ |
| Epithelium | $\gamma\delta$ T Cells | CD45+ EpCAM- TCR$\gamma\delta$+ |
| Epithelium | CD8$\alpha\alpha$+ TCR$\alpha\beta$ Natural IEL | CD45+ EpCAM- TCRβ+ CD8α+ CD8β- |
| Epithelium | CD8$\alpha\alpha$+ TCR$\gamma\delta$ Natural IEL | CD45+ EpCAM- TCR$\gamma\delta$+ CD4- CD8β- CD8α+ |
| Epithelium | CD8$\alpha\beta$+ TCR$\alpha\beta$ Induced IEL | CD45+ EpCAM- TCRβ+ CD8α+ CD8β+ |
| Lamina Propria/Epithelium | Eosinophils | CD45+ CD11b+ Ly6G- SiglecF+ SSC high |
| Lamina Propria/Epithelium | Monocytes | CD45+ CD11b+ Ly6C+ Ly6G- |
| Lamina Propria/Epithelium | Neutrophils | CD45+ CD11b+ Ly6C+/- Ly6G+ |
| Lamina Propria/Epithelium | Macrophages | CD45+ CD11b+ F4/80+ |
| Lamina Propria | $\alpha\beta$ T Cells | TCRβ+ |
| Lamina Propria | $\gamma\delta$ T Cells | TCR$\gamma\delta$+ |
| Lamina Propria | CD4+ T Cells | TCRβ+ CD4+ CD8- |
| Lamina Propria | CD8+ T cells | TCRβ+ CD4- CD8+ |
| Lamina Propria | Roryt+ | TCRβ+ CD4+ CD8- Roryt+ |
| Lamina Propria | Treg | TCRβ+ CD4+ CD8- FoxP3+ |
| Lamina Propria | Roryt+ Treg | TCRβ+ CD4+ CD8- FoxP3+ Roryt+ |
| Lamina Propria | CD44+ T cells | TCRβ+ CD44+ |
| Lamina Propria | B Cells | B220+ CD19+ |
| Lamina Propria | IgA+ Plasma Cells | Lin- (CD3-TCRβ-CD4-CD11c-NK1.1-F4/80-) B220- IgA+ |

| Target | Fluorophore | Clone | Vendor | Catalog # |
| --- | --- | --- | --- | --- |
| B220 | PE-Cy7 | RA3-6B2 | Biolegend | 103222 |
| CD11b | PerCP-Cy5.5 | M1/70 | Biolegend | 101228 |
| CD11c | Brilliant Violet 421 | N418 | Biolegend | 117330 |
| CD19 | Brilliant Violet 421 | 6D5 | Biolegend | 115538 |
| CD4 | FITC | RM4-5 | Biolegend | 100510 |
| CD4 | APC | RM4-5 | Biolegend | 100516 |
| CD44 | APC-Cy7 | IM7 | Biolegend | 103028 |
| CD45 | APC eFluor780 | 30-F11 | eBioscience/Invitrogen | 47-0451-82 |
| CD8α | PE-Cy7 | 53-6.7 | Biolegend | 100722 |
| CD8β | FITC | YTS156.7.7 | Biolegend | 126605 |
| EpCAM | Brilliant Violet 421 | G8.8 | Biolegend | 118225 |
| F4/80 | APC | BM8 | Biolegend | 123116 |
| FoxP3 | APC | FJK-16s | eBioscience/Invitrogen | 17-5773-82 |
| Granzyme A | PE | 3G8.5 | Biolegend | 149704 |
| IgA | PE | mA-6E1 | eBioscience/Invitrogen | 12-4204-83 |
| LY6C | FITC | AL-21 | BD Biosciences | 553104 |
| LY6G | PE-Cy7 | 1A8 | Biolegend | 127618 |
| PD1 | APC-Cy7 | 29F.1A12 | Biolegend | 135224 |
| Roryt | PE | B2D | eBioscience/Invitrogen | 12-6981-82 |
| SiglecF | PE | E50-2440 | BD Biosciences | 552128 |
| TCRβ | PerCP-Cy5.5 | H57-597 | Biolegend | 109228 |
| TCR$\gamma\delta$ | Brilliant Violet 421 | GL3 | Biolegend | 118120 |
| TCR$\gamma\delta$ | PE | GL3 | Biolegend | 118108 |
| CD3 | PerCP-Cy5.5 | 145-2C11 | Biolegend | 1000328 |
| CD4 | PerCP-Cy5.5 | RM4-5 | Biolegend | 100539 |
| CD11c | PerCP-Cy5.5 | N418 | Biolegend | 117327 |
| NK1.1 | PerCP-Cy5.5 | PK136 | Biolegend | 108728 |
| F4/80 | PerCP-Cy5.5 | BM8 | Biolegend | 123127 |
| Live dead | Aqua | -- | Biolegend | 423101 |

### Protein Quantification

Ileum samples were homogenized in Lysing Matrix F (MP Biomedicals, ref. 6540440) and 500uL of 1x HALT (Thermo Scientific, ref. 78429) in T-PER (Thermo Scientific, ref. 78510). Homogenates were centrifuged at 4 °C and 10,000 xg for 5 minutes. Protein concentration was determined by BCA assay (Thermo Scientific, ref. 23227) using the manufacturer’s protocol. Lysates were normalized and stored at −80 °C until analysis. Samples were sent to UVA’s Flow Cytometry Core Facility where the Luminex assay was performed according to the manufacturer’s protocol, which is as follows: 200uL of Wash Buffer (room-temperature 1x Wash Buffer in deionized water) was added to each well of a clean plate and sealed. Plate was mixed for 10 minutes on a plate shaker at 20-25 °C. After removing the wash buffer, 25uL of standard or control was added to the corresponding wells. 25uL of assay buffer were added to background and sample wells. 25uL of previously mentioned lysis buffer was added to the background, standards, and control wells. After vortexing the mixing bottle, 25uL of the mixed or premixed beads were added to each well. The plate was sealed, covered with foil, and incubated overnight at 2-8 °C with agitation. After removing the well contents and washing the plate twice, 25uL of detection antibody was added to each well. The plate was sealed, covered with foil, and incubated for 1 hour at 20-25 °C with agitation. 25uL of Streptavidin-Phycoerythrin was added to the wells containing detection antibody. The plate was sealed, covered with foil, and incubated for 30 minutes at 20-25 °C with agitation. After removing the well contents and washing the plate twice, 150uL of Sheath Fluid PLUS was added to all wells. The beads were resuspended on a plate shaker for 5 minutes and the plate was measured on the Luminex xMAP Intelliflex. Sample analyte concentration was calculated by fitting the median fluorescence intensity (MFI) data to that of a standard curve, validated by lot-matched quality controls using the Milliplex Analyst software.

CCL5 and IL-1β were quantified by ELISA assay (R&D Systems) on the tissue lysates prepared as described above according to the manufacturer’s recommendation. Briefly, the capture antibody was coated onto a 96-well half-area plate (Corning) in PBS overnight at room temperature. On the following day, the plates were washed three times with 200 uL of Wash Buffer (0.05% Tween 20 in PBS) and blocked for one hour. After blocking and incubation, plates were washed again before adding samples of small intestinal tissue lysate as well as a standard curve. The plate was incubated for 2 hours at room temperature before being washed as described above. Detection Antibody provided in the kit was added at the suggested concentration and incubated for another 2 hours at room temperature. After incubation, plates were washed and Substrate Solution (1:1 mixture of Color Reagent A (H_2_O_2_) and Color Reagent B (Tetramethylbenzidine)) was added to each well and incubated at room temperature for 20 minutes avoiding direct light. After incubation, Stop Solution (2 N H_2_SO_4_) was added to each well. Plates were reader an optical density of 450 with background subtraction at 570 nm using a Tecan plate reader.

## QUANTIFICATION AND STATISTICAL ANALYSIS

Statistical analyses were performed in GraphPad Prism unless otherwise noted. Statistical details including the number of animals or samples can be found in figure legends. Statistical significance was assessed by Mann-Whitney U test when comparing between two groups, or by Two-Way ANOVA with Šídák’s multiple comparisons test when comparing >2 groups. P values are shown in figures or tables for samples with significant differences.

